# Phase-specific manipulation of neural oscillatory activity by transcranial alternating current stimulation

**DOI:** 10.1101/579631

**Authors:** Marina Fiene, Bettina C. Schwab, Jonas Misselhorn, Christoph S. Herrmann, Till R. Schneider, Andreas K. Engel

**Affiliations:** Department of Neurophysiology and Pathophysiology, University Medical Center Hamburg-Eppendorf, Hamburg, 20246 Germany; Experimental Psychology Lab, Department of Psychology, Cluster of Excellence “Hearing4all”, European Medical School, Carl von Ossietzky University Oldenburg, Oldenburg, 26129 Germany; Research Center Neurosensory Science, Carl von Ossietzky University Oldenburg, Oldenburg, 26129 Germany

**Author notes:** **Corresponding author** Marina Fiene, Department of Neurophysiology and Pathophysiology, University Medical Center Hamburg-Eppendorf, Martinistraße 52, 20246 Hamburg, Germany.

**Keywords:** Transcranial alternating current stimulation, Electroencephalogram, Entrainment, Alpha oscillations, Visual flicker

## Abstract

**Background:** Oscillatory phase has been proposed as a key parameter defining the spatiotemporal structure of neural activity. To enhance our understanding of brain rhythms and improve clinical outcomes in pathological conditions, phase-specific modulation of oscillations by transcranial alternating current stimulation (tACS) emerged as a promising approach. However, the effectiveness of tACS in humans is still critically debated.

**Objective:** Here, we investigated the phase-specificity of tACS effects on visually evoked steady state responses (SSRs) in 24 healthy human participants of either sex.

**Methods:** To this end, we used an intermittent electrical stimulation protocol and assessed the influence of tACS on SSR amplitude in the interval immediately following tACS.

**Results:** We observed that the phase shift between flicker and tACS modulates evoked SSR amplitudes. The tACS effect size was dependent on the strength of flicker-evoked oscillatory activity, with larger effects in participants showing weaker locking of neural responses to flicker phase. Neural sources of phase-specific effects were localized in the parieto-occipital cortex within flicker-entrained regions. Importantly, the optimal phase shift between flicker and tACS associated with strongest SSRs was correlated with cortical SSR onset delays over the visual cortex.

**Conclusions:** Overall, our data provide electrophysiological evidence for phase-specific modulations of oscillatory activity by tACS in humans. As the optimal timing of tACS application was dependent on neural conduction times as measured by SSR onset delays, data suggest that the interaction between tACS effect and SSR was cortical in nature. These findings corroborate the physiological efficacy of tACS and highlight its potential for controlled modulations of brain signals.

## Introduction

Oscillations are a prominent feature of activity in neural networks proposed to be of particular importance for neural processing and, by extension, for normal and pathological brain function [1–4]. Modulation of such cortical activity by means of non-invasive brain stimulation represents a promising approach to advance knowledge on the functional role of the temporal patterning of neural activity and to improve clinical outcomes in disorders related to aberrant neural synchrony. Specifically, transcranial alternating current stimulation (tACS) is widely applied with the aim to entrain neural activity, implying a periodic modulation of membrane potentials and a phase alignment of intrinsic oscillations to the tACS phase. This assumption is supported by in vitro slice preparations and recordings in primates showing phase-specific modulations of neuronal spiking patterns by externally applied electric fields [5–8]. However, evidence from invasive recordings in animals may not directly be transferable to tACS effects in the human brain. Despite its broad use in basic and clinical science, the neural mechanism of tACS in humans is still under investigation and its overall efficacy has recently been challenged. Measurements of the electric field strength in human cadaver brains and epilepsy patients with implanted electrodes showed that most of the scalp-applied current is attenuated by skin and skull thereby questioning significant neural entrainment [9–11]. These findings, paralleled by inconsistent behavioral tACS effects [12–14], encouraged a reconsideration of study requirements to investigate the physiological mechanisms of tACS.

The theoretically ideal approach to directly measure tACS-induced effects on neural activity is complicated by strong, hard-to-predict electrical artifacts in concurrent recordings of neural activity [15,16]. To provide evidence for online phase-specific neuronal effects despite these methodological constraints, most studies assessed tACS effects on indirect measures assumed to be mediated by brain rhythms. For instance, tACS applied in the alpha and beta range was shown to lead to cyclic modulations of auditory, somatosensory or visual stimulus perception [17–20]. Further insight has been provided in the motor domain, by showing tACS-phase-dependent changes in cortical excitability [21–23] or peripheral tremor [24–26]. Yet, tACS-effects on the motor cortex were shown to be dominated by cutaneous co-stimulation of peripheral nerves in the skin leading to rhythmic activation of the somatosensory system rather than by transcranial modulation of cortical tissue [27]. Related evidence might thus not necessarily be generalizable to transcranial entrainment effects in cortical regions apart from the motor cortex. Further, prior conclusions only hold true under the assumption that the chosen outcome measures, e.g., behavior or cortical excitability, are reliable proxies for cortical oscillatory activity.

Here, we used an innovative approach that combines rhythmic electrical with rhythmic sensory stimulation to target the phase-specificity of tACS effects directly on cortical oscillatory activity. Via electroencephalography (EEG), we measured neural tACS effects on precisely controlled brain rhythms evoked by visual flicker, i.e., visually evoked steady state responses (SSRs), as a function of six phase shifts between tACS and flicker onset. Using an intermittent electrical stimulation protocol, concurrent 10 Hz flicker and multi-electrode tACS targeting the occipital cortex were applied in two sessions under active and sham tACS. Artifact free epochs in the immediate interval following tACS were used for SSR amplitude analysis (Figure 1). We predicted that the phase shift between flicker- and tACS-induced membrane voltage fluctuations should systematically modulate the amplitude of net SSRs.

**Figure 1.**
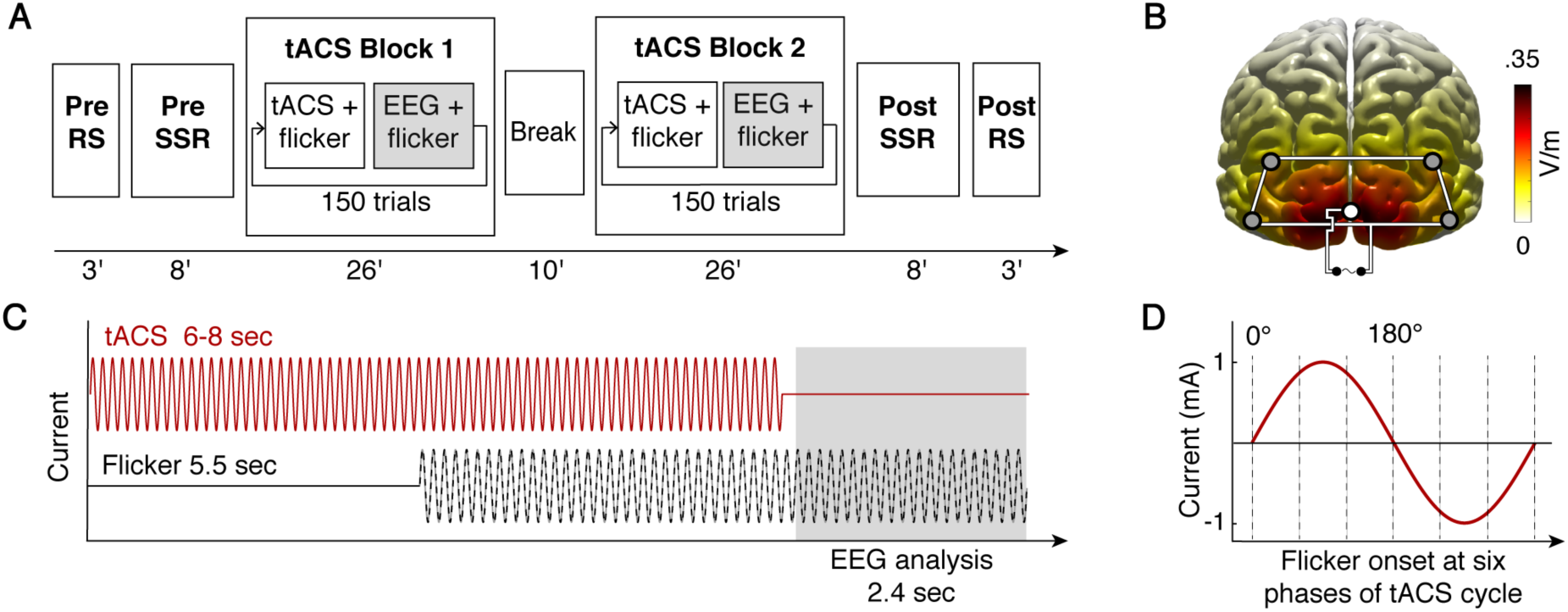
Experimental setup. (A) Schematic display of the experimental timeline. The experiment was conducted on two separate days for active tACS and sham stimulation. Resting state (RS) and visual evoked steady state response (SSR) data without tACS were recorded pre and post to the tACS blocks. (B) tACS electrode montage and power of the tACS-induced stimulation field. The occipital electrode montage induced highest current densities over the primary visual cortex and current spread in surrounding areas. Maximum peak-to-peak difference in field strength is about 0.7 V/m. (C) Illustration of one trial of the intermittent tACS protocol. Each trial started with tACS only, followed by flicker onset after 3-5 sec. The flicker continued for 5.5 sec until the end of the trial. The gray-shaded tACS free intervals were used for EEG data analysis. (D) To vary the phase between neural activity induced by electrical and sensory rhythmic stimulation, visual flicker started at six different phase shifts relative to the tACS cycle. 50 trials per phase angle were presented in randomized order.

## Materials and Methods

### Participants

24 healthy volunteers (mean age 25.1 ± 3.3 years; 16 female; 8 male; 22 right-handed) were recruited from the University Medical Center Hamburg-Eppendorf, Germany. The sample size was chosen based on the effect sizes between sham and active tACS in previous studies showing phase-dependent stimulation effects on sensory perception or peripheral tremor [18,24,28]. All participants reported normal or corrected-to-normal vision, no history of psychiatric or neurological disorders and were blind towards the tACS stimulation sequence. The experimental protocol was approved by the ethics committee of the Hamburg Medical Association and was conducted in accordance with the Declaration of Helsinki. All participants gave written informed consent before participation. Participants were monetarily compensated and debriefed immediately after the experiment.

### Experimental design

All subjects participated in two separate sessions in a randomized, counterbalanced order once under sham and active tACS. Participants were seated in a dimly lit, electrically shielded and sound-attenuated EEG chamber at a distance of approximately 40 cm to a white light emitting diode (LED). Due to a nonlinear relation between forward voltage and luminosity of the used LED, we assessed LED luminosity per duty cycle of the pulse-width modulation by a luminance meter (LS-100, Konica Minolta). The resulting function was multiplied with a pure 10 Hz sine to generate a 10 Hz luminance signal. The LED had a maximal light intensity of 100 cd/m^2^. Prior to the main tACS block, 3 min eyes-open resting state EEG (Pre RS) was recorded while participants fixated the white illuminated LED. Afterwards, EEG during an 8 min SSR block without tACS (Pre SSR) was recorded. Single flicker periods (50 in total) were presented for 5.5 sec followed by short breaks of 3-5 sec. The timing of flicker presentation was identical to the procedure used in the main tACS block. In the tACS block, EEG was recorded continuously during the intermittent tACS protocol. Each trial started with tACS only and was followed by flicker onset after 3-5 sec. The flicker continued until the end of the trial for 5.5 sec in total. tACS free intervals of 2.5 sec were used for EEG data analysis. Single tACS periods had a duration of 6-8 sec. Between successive tACS epochs, tACS phase was continuous by adjusting only the amplitude but not the phase of the sine wave spanning the whole testing session. To account for the varying phase delay between neural activity induced by sensory and electrical stimulation, the visual flicker started at six different phase angles (0°, 60°, 120°, 180°, 240° and 300°) relative to the tACS cycle. 50 trials were recorded for each of the six phase shift conditions (300 trials in total). The sequence of trials was randomized such that every phase shift condition was followed equally likely by all other conditions. Moreover, to avoid confounding time-on-task effects on alpha amplitude, trials belonging to different conditions were evenly distributed over the course of the testing session by iterate permutation. The tACS block was divided in two blocks interrupted by a 10 min break. After the tACS block, another SSR block without tACS (Post SSR) and resting state EEG (Post RS) were recorded.

To ensure that participants fixate the LED and thus to enhance SSR detection, participants were instructed to detect oddball trials in which LED luminance changed after 3-4 sec. Participants responded per button press as soon as they detected a luminance change. 16 oddball trials were included in the tACS block and two oddball trials in the pre and post SSR blocks, respectively. Participants showed a mean detection accuracy of 97.66 ± 9.99 % (mean ± sd) during the tACS blocks and 97.92 ± 12.39 % during the pre and post SSR blocks.

### Electrophysiological recording

EEG was recorded continuously from 64 Ag/AgCl electrodes (12 mm diameter) mounted in an elastic cap (Easycap). The electrooculogram (EOG) was recorded from two electrodes placed below both eyes with the EEG referenced to the nose tip. Electrodes were prepared with abrasive conducting gel (Abralyt 2000, Easycap) keeping impedance below 20 kΩ. EEG data were recorded using BrainAmp DC amplifiers (Brain Products GmbH) and the corresponding software (Brain Products GmbH, Recorder 1.20). Data were recorded with an online passband of 0.016-250 Hz and digitized with a sampling rate of 1000 Hz.

### tACS and electric field simulation

Multi-electrode tACS was applied via five additional Ag/AgCl electrodes (12 mm diameter) mounted between EEG electrodes. The montage was connected to a battery-driven current source (DC-Stimulator Plus, NeuroConn). After preparation with Signa electrolyte gel (Parker Laboratories Inc.), impedance of each outer electrode to the middle electrode was kept below 20 kΩ. Moreover, impedances were kept comparable to achieve an evenly distributed electric field. A sinusoidal alternating current of 2 mA peak-to-peak was applied at 10 Hz using an intermittent stimulation protocol. It has previously been shown that flicker-evoked SSRs show strongest amplitudes in the alpha frequency range [29]. Also, most reliable evidence for aftereffects of tACS on oscillatory power has been demonstrated in the alpha band that was proposed to result from online neural phase-alignment [30,31]. Thus, we chose 10 Hz to test for the general feasibility of phase-specific effects by tACS, although our approach is not limited to this stimulation frequency. During sham and active stimulation, the current was ramped in over 15 sec to 2 mA. After 90 sec of continuous stimulation, tACS periods of 6-8 sec without ramp were applied interrupted by short breaks of 2.5 sec for EEG analysis. Total stimulation time during the active session was 40 min. For sham stimulation, the current was ramped in and out over 15 sec each. This procedure ensured that in both stimulation conditions, participants experienced initial itching sensation. All participants confirmed that stimulation was acceptable and did not induce painful skin sensations. The most common side-effect was a tingling feeling on the scalp under the electrode. During the debriefing at the end of the final session, except for two subjects all participants correctly rated the sequence of active and sham stimulation. Further, exploration of the experience of phosphenes during tACS in the beginning of each session revealed that no participant reported phosphene perception. This finding is in line with our simulation of the tACS-induced electric field which greatly diminishes with increasing distance to stimulation electrodes and is expected to be low in the retina [32].

Placement of tACS electrodes was chosen to target brain regions activated by visual stimulation. We aimed to achieve largest magnitudes of current density in the primary visual cortex to maximally interfere with early stages of cortical sensory processing. Quantitative field distributions 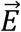 could directly be computed by linear superposition of lead fields 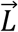, weighted by the injected currents *α*_*i*_ at electrodes *i* = {1,2,3,4,5}:

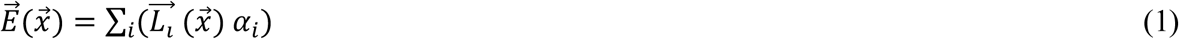

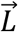 was estimated with the boundary element method using a volume conduction model by Oostenveld et al. [33] at 5 mm spatial resolution.

## Data analysis

### Data preprocessing

64-channel EEG data were preprocessed and analyzed in Matlab (The MathWorks Inc.) using the analysis toolbox FieldTrip [34]. Circular statistics were calculated with the CircStat toolbox [35]. EEG recordings during the pre and post SSR blocks without tACS were segmented into epochs from −1.5 to 6.5 sec around flicker onset. For the tACS block, EEG was segmented into epochs from 0.1 to 2.5 sec after tACS offset. The first 0.1 sec after stimulation offset were rejected before filtering to avoid leakage of capacitive discharge effects after tACS offset, appearing as a short-lasting voltage drift back to baseline EEG level [30,36]. Examination of voltage across 20 ms time segments confirmed no further significant voltage offset change beyond 0.1 sec after the end of stimulation. EEG data were bandpass filtered from 1 to 20 Hz using a Hamming-windowed sinc FIR filter. Oddball trials and trials during which participants erroneously responded via button press were excluded. To automatically screen for high noise level electrodes, channels whose standard deviation exceeded 3 times the median standard deviation of all channels per participant were excluded. On average, 4.0 ± 1.9 channels were removed. Data were downsampled to 100 Hz. Afterwards, independent component analysis (ICA) was computed using the infomax ICA algorithm [37]. Remaining eye-movement, cardiac and noise artifacts were removed based on visual inspection of the components’ time course, spectrum and topography. For eye-blink artifact removal, Pearson linear correlation coefficient between EOG and ICA components was calculated and components showing correlation coefficients greater than .25 were rejected. On average, 20.1 ± 3.9 components were excluded. Finally, all trials were visually inspected and trials containing artifacts that had not been detected by previous steps were manually removed. On average, 49.19 ± 1.2 out of 50 trials remained for the pre and post SSR blocks and 296.9 ± 3.5 out of 300 trials remained for the tACS block per participant.

### Source reconstruction

At the cortical source level, neural activity was estimated using exact low resolution brain electromagnetic tomography (eLORETA). Before source projection, data were bandpass filtered between 7-13 Hz with a 4^th^ order butterworth filter. The cortical grid was defined using the Automated Anatomical Labeling (AAL) atlas consisting of 38 regions after excluding subcortical and cerebellar structures [38]. Three-dimensional time series were estimated in a linearly spaced grid of 4077 cortical points with 7.5 mm distance in all three spatial directions using the boundary element method volume conduction model by Oostenveld et al. [33]. eLORETA was applied per trial with 5% regularization. To reduce spatial dimensions, the three resulting time series were projected to the direction of maximal power at each grid point.

### Spectral analysis and SSR amplitude modulation indices

EEG spectral analysis was performed on sensor and source level. We expected to find phase-specific tACS-induced modulations of SSR amplitude at electrode positions showing a clear response to visual flicker, i.e., where phase alignment to flicker stimulation is prominent. Therefore, we assessed the spatial distribution of phase locking of EEG to visual flicker in the pre SSR blocks without tACS for the sham and active tACS testing session. A Hilbert transform was computed on the flicker signal and EEG time series during flicker stimulation (1.0 to 5.5 sec after flicker onset) for each trial at each electrode and source level grid point. The first 1 sec after flicker onset was removed to ensure the SSR has fully build up. To assess phase stability, the time-dependent differences in phase signals (*ϕ*_*flicker*_ (*t*) − *ϕ*_*EEG*_ (*t*)) were computed and the extent of entrainment quantified by the phase locking value (PLV) over all trials.

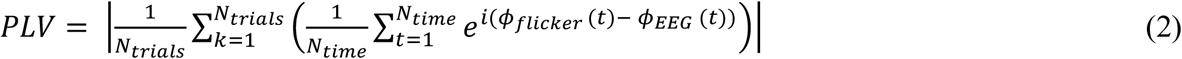

For sensor level analysis, we selected those channels showing prominent phase entrainment with mean PLV > 0.5 over all participants. Pearson correlation coefficient was computed to quantify the relation between PLV and SSR amplitude increase from resting state to pre SSR blocks. SSR amplitude was calculated by Fourier transform of 1 sec time windows with an overlap of 0.5 sec for each 3 min resting state block (pre RS) and for each 4.5 sec trial of the pre SSR blocks. To correct for global alpha amplitude fluctuations over time independent of experimental manipulation, SSR amplitude was defined as a relative amplitude value by subtracting the mean amplitude at neighboring frequencies (9 and 11 Hz) from the absolute amplitude at the stimulation frequency.

For analysis of SSR amplitude modulation during the tACS block, we computed the Fourier transform of each trial separately for the time epochs of 0.1 to 1.1 sec, 0.6 to 1.6 sec and 1.1 to 2.1 sec after tACS offset. To assess modulation of SSR amplitude dependent on the relative phase shift between tACS and flicker onset, bar diagrams were constructed showing the variation in SSR amplitude over phase shift conditions. SSR amplitude was normalized by the sum of amplitude values over all six conditions. Assignment of trials to phase conditions for the sham session was performed using the same trial numbers as under tACS stimulation, thereby ensuring equal time-on-task effects for both sessions. If there was no phase-specific amplitude modulation by concurrent tACS, bar graphs should be uniformly distributed. In case of phase-specific interactions, the SSR amplitude should be higher in a condition that shows an optimal phase shift for that participant. Assuming that the optimal phase shift between flicker and tACS onset leading to increased SSR amplitudes might vary between participants, any preference in phase shift condition compared to sham can be considered as evidence for phase specific tACS effects. Amplitude modulation was quantified by three modulation measures. First, we calculated the Kullback-Leibler divergence (D_KL_) which is a common premetric used in statistics to calculate the difference between two distributions. The deviation of the observed amplitude distribution *P* from the uniform distribution *U* is defined as

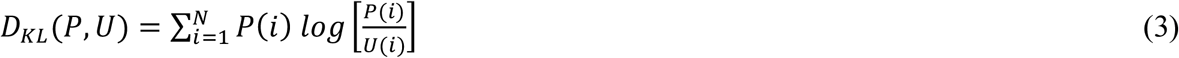

where *N* is the number of phase bins [39]. Strict uniformly distributed amplitude values would be reflected in D_KL_ = 0 whereas D_KL_ increases the further away *P* gets from *U*. While D_KL_ measures the overall deviation of amplitude values from uniform distribution, we additionally assessed whether the tACS-induced amplitude modulation over phase conditions follows a cyclic pattern, as hypothesized based on the sinusoidal nature of the tACS signal. Therefore, we calculated the phase locking value of amplitude values to phase condition. This cyclic amplitude phase locking (APL) index is defined as

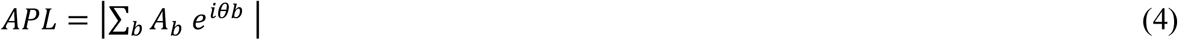

where *A* and *θ* are the normalized relative amplitude and phase shift for each of the six phase shift conditions *b*. To further investigate whether the tACS-induced SSR amplitude modulation can be described by a sinusoidal pattern, we fitted one-cycle sine waves to SSR amplitude bar diagram values [18,40]. The following equation was fit to the six normalized SSR amplitude values per participant and stimulation condition,

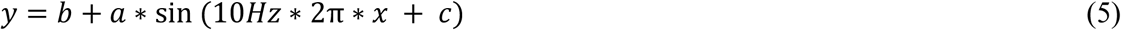

where *x* corresponds to the flicker-tACS phase shifts during one cycle (*x* = {0, 1/6T, 2/6T,… 5/6*T} with period T = 0.1 sec). The parameters *a, b* and *c* were estimated by the fitting procedure where *b* is the intercept, *a* the amplitude and *c* the phase shift. Parameters *a* and *b* were bound between −1 and 1 and *c* between 0 and 2π. Start values were drawn randomly from within the range of restrictions for each parameter. Model fits were computed using the Matlab fittype function allowing for 1000 iterations to find the best model. We examined the absolute sine fit amplitude values as index for the strength of SSR amplitude modulation. The significance of the sine fit was assessed by dependent samples t-tests comparing the proportion of explained variance R^2^ as goodness-of-fit index between sham and active tACS session.

### Statistical analysis

For each of the three modulation indices, we calculated dependent samples t-tests for the epochs of 0.1 to 1.1 sec, 0.6 to 1.6 sec and 1.1 to 2.1 sec after tACS offset between sham and active tACS on sensor level. Effect sizes were determined by calculating Cohen’s d_z_ for paired t-test comparisons. To account for comparisons in multiple time windows, we applied Bonferroni correction to the significance level and report test statistics if the corresponding *p*-value fell below the corrected alpha of α_bonf_ = .0167 (= .05/3).

For further investigation of tACS effects, we analyzed the APL parameter resulting from the epoch immediately following tACS offset in source space. APL is the more specific, phase-dependent modulation index relative to D_KL_ and a more robust measure not requiring any presuppositions on parameter ranges as compared to sine fit. For source localization of APL effects, dependent samples t-tests were computed for each grid point using cluster-based permutation tests to correct for multiple comparisons. Clusters were obtained by summing up t-values which were adjacent in space and below an alpha level of 5 % (two-sided). A permutation distribution was generated by randomly shuffling APL values between active and sham condition within participants in each of 1000 iterations. The observed cluster was considered statistically significant when the sum of t-values exceeded 95 % of the permutation distribution. To further examine whether the strength of tACS effects quantified by APL is dependent on the underlying strength of flicker-evoked oscillatory activity, we assessed Pearson correlation coefficients between APL and EEG-flicker PLV for sham and active tACS blocks. Moreover, we assessed the dependency of the increase in APL from sham to tACS on the change in mean SSR amplitude, the standard deviation of SSR amplitudes or EEG-flicker PLVs over all trials per participant by computing Pearson correlation coefficients.

Finally, to disentangle retinal from cortical stimulation effects, we examined the dependency of optimal flicker-tACS phase shifts on the cortical flicker-SSR phase delays by computing circular-circular correlations per channel using cluster-based permutation tests [35]. The flicker-SSR phase delay was computed as the mean time-dependent difference in phase signals (*ϕ*_*flicker*_(*t*) − *ϕ*_*EEG*_(*t*)) based on the Hilbert transform of the EEG and flicker signal over all trials of the tACS block. Cluster-level test statistics were calculated by taking the maximum average of correlation coefficients across neighboring electrodes showing significance of circular correlations (p < .05). The permutation distribution was generated by between-subject randomization of flicker-tACS and SSR onset phase delay data over 1000 iterations.

## Results

### Flicker-evoked phase alignment in the parieto-occipital cortex

To analyze evoked flicker entrainment, we assessed the degree and spatial distribution of PLVs between EEG and 10 Hz flicker stimulation. Estimated time series on source level displayed a prominent peak of parieto-occipital EEG-flicker PLV (Figure 2B). Strong phase locking was located in the superior and middle occipital gyrus as well as in bilateral precuneus and cuneus. For sensor level analysis, four parieto-occipital channels showing a population mean PLV > 0.5 (corresponding to electrodes O1, O2, POz, Pz) were selected for further analysis of SSR amplitude modulation by tACS. Mean PLV for the selected channels during both pre SSR blocks over all participants was 0.56 ± 0.16 (mean ± sd). As illustrated in the lower part of Figure 2B, data show varying phase delays between flicker onset and cortical SSR across participants related to differences in neural conduction delays along the visual pathway. Further, amplitude spectra revealed an amplitude increase at the flicker stimulation frequency compared to resting state (Figure 2C). Between-subject correlation showed that the strength of EEG-flicker PLV was positively correlated with the increase in SSR amplitude from resting state to pre SSR blocks (*r* = .43, *p* = .037) (Figure 2D).

**Figure 2.**
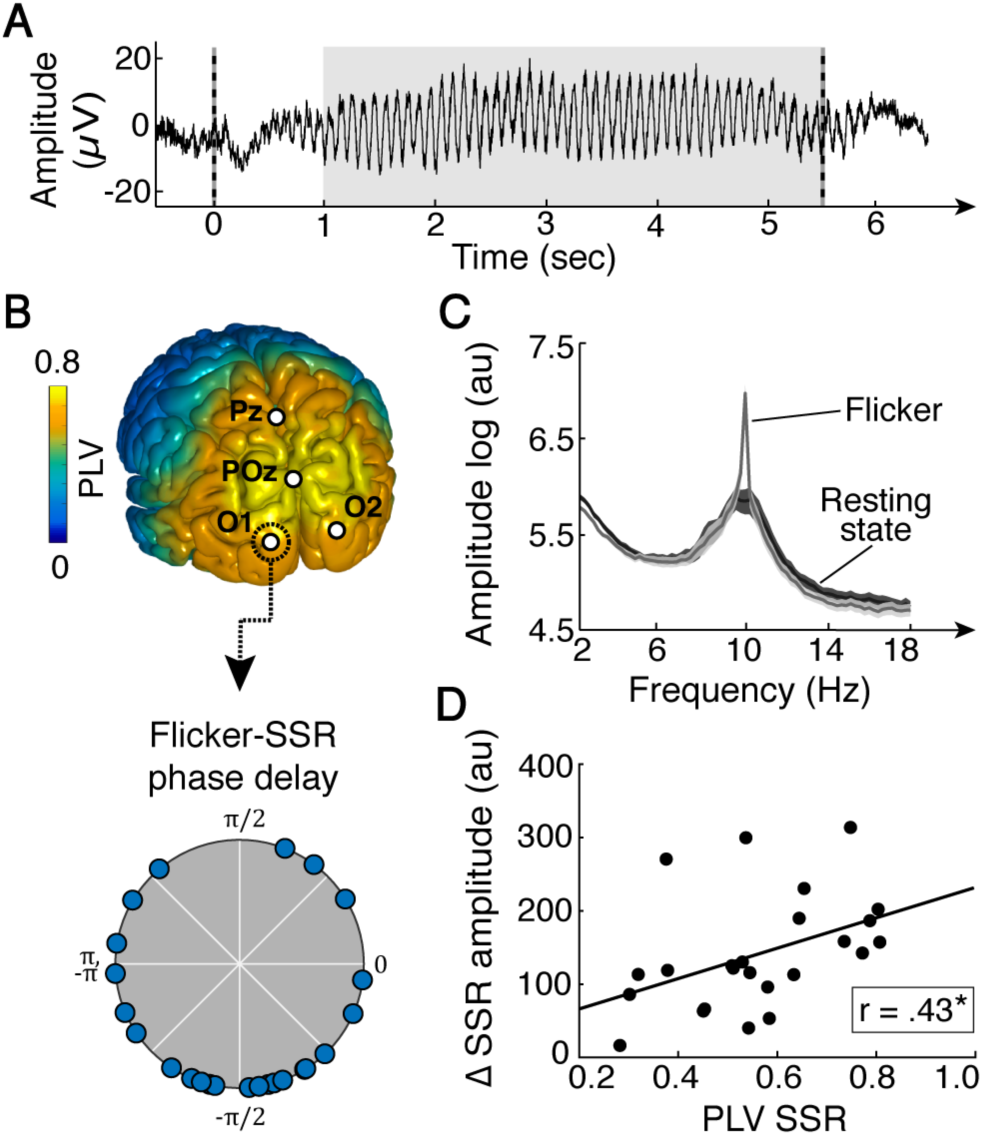
Phase locking values (PLVs) of steady state responses (SSRs) to flicker phase over all trials of the active and sham pre SSR blocks. (A) SSR during 5.5 sec flicker stimulation computed as the average of 50 raw EEG traces of one exemplary participant at channel POz. Dashed lines mark the onset and offset of visual flicker. The gray-shaded area depicts the time window used for EEG data analysis. (B) The upper part depicts the spatial distribution of mean PLVs over all participants with maximal phase locking over the parieto-occipital cortex. Markers correspond to channels selected on sensor level for SSR amplitude analysis. The lower part shows the interindividual variation in phase delays between flicker onset and cortical SSR exemplary for electrode O1. (C) Mean amplitude spectrum during flicker stimulation and resting state over all participants for the four selected channels on sensor level. Shaded areas indicate the standard error. (D) The strength of EEG-flicker PLV at the four parieto-occipital channels is positively correlated with the increase in SSR amplitude from resting state to flicker stimulation during pre SSR blocks across participants (*r* = .43, *p* = .037, n = 24). * *p* < .05.

### Phase-dependent SSR amplitude modulation by tACS

To test the hypothesis that simultaneous tACS and flicker stimulation lead to phase-specific interactions between evoked neural activity and electric field, we assessed the degree of tACS-phase-dependence of SSR amplitude values via three modulation measures: (1) the deviation of amplitude values from a uniform distribution (D_KL_), (2) the locking of amplitude values to phase conditions (APL) and (3) one-cycle sine fits to SSR amplitude values. Figure 3A shows normalized SSR amplitude bar diagrams over the six phase shift conditions for one exemplary participant. During sham, data are almost uniformly distributed across phase bins, showing small fluctuations in SSR amplitude with no preference for a particular phase shift condition. On the contrary, the bar diagram for active stimulation shows phase-dependent modulations of SSR amplitudes indicating an interaction between tACS-induced electric field and flicker-evoked oscillatory activity. On sensor level, analysis of the deviation of amplitude values from uniform distribution revealed a significantly greater D_KL_ under tACS compared to sham within the first 1 sec epoch (0.1 to 1.1 sec) after tACS offset (*t*(23) = 3.21, *p* = .004 < α_bonf_, *d*_*z*_ = 0.66) (Figure 3B). Phase locking of amplitude values to phase condition measured by APL was significantly larger after tACS (*t*(23) = 3.12, *p* = .005 < α_bonf_, *d*_*z*_ = 0.64). In accordance, also the one-cycle sine wave fit to SSR amplitudes showed a significantly greater amplitude of the sine fit under tACS (*t*(23) = 3.52, *p* = .002 < α_bonf_, *d*_*z*_ = 0.72) (Figure 3B). The proportion of explained variance R^2^ by sine fit was greater for SSR amplitude modulation under tACS compared to sham (*t*(23) = 2.65, *p* = .014, *d*_*z*_ = 0.54). In successive time windows, the modulatory tACS effect decayed for all three modulation parameters ([0.6, 1.6] sec: D_KL_, *t*(23) = 0.46, *p* = .650, *d*_*z*_ = 0.09; APL, *t*(23) = 2.29, *p* = .032, *d*_*z*_ = 0.47; Sine fit, *t*(23) = 2.23, *p* = .036, *d*_*z*_ = 0.45; [1.1, 2.1] sec: D_KL_, *t*(23) = −0.30, *p* = .769, *d*_*z*_ = −0.06; APL, *t*(23) = −0.25, *p* = .806, *d*_*z*_ = −0.05; Sine fit, *t*(23) = 0.52, *p* = .610, *d*_*z*_ = 0.11; Figure 3E). Importantly, the phase-dependent effects of tACS on SSR amplitude were not caused by overall 10 Hz alpha amplitude differences between conditions (Figure 3C). A repeated-measures t-test revealed no significant difference in mean SSR amplitude (*t*(23) = −1.02, *p* = .317, *d*_*z*_ = 0.21) over all trials per participant between tACS and sham condition during the first 1 sec interval after tACS offset (Figure 3D). Also, the standard deviation of SSR amplitude values over all trials per participant did not differ between sham and active stimulation condition (*t*(23) = -.31, *p* = .759, *d*_*z*_ = 0.06). This implies a strengthening or dampening of SSR amplitudes dependent on the phase shift between flicker and tACS stimulation.

**Figure 3.**
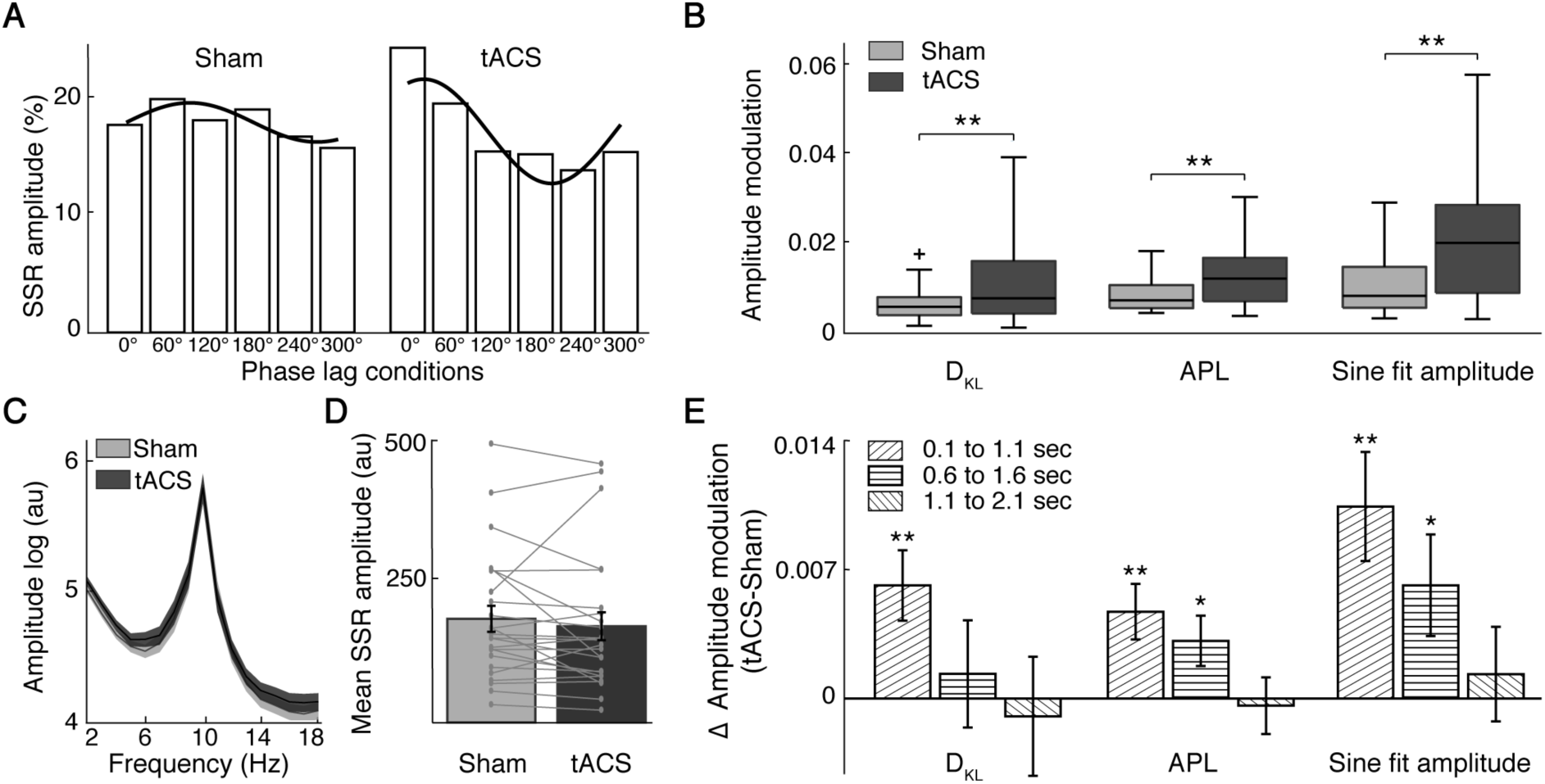
Steady state response (SSR) amplitude modulation by tACS shown for the time window of 0.1 to 1.1 sec after stimulation offset during the tACS block on sensor level. (A) SSR amplitude bar graphs for one exemplary participant. SSR amplitude values differ in the degree of phase dependence under tACS compared to sham. Black lines show sinusoidal fits to the data. (B) Significantly increased Kullback-Leibler divergence (D_KL_), enhanced phase locking of amplitude values to phase condition (APL) and sine fit amplitudes under tACS relative to sham. (C, D) Phase-dependent amplitude modulation is not paralleled by differences in mean absolute 10 Hz alpha amplitude or mean SSR amplitude over all trials and participants between stimulation conditions. Shaded areas and error bars indicate the standard error. (E) Decay in tACS effect size over three time windows after tACS offset. Bar graphs show the mean difference in SSR amplitude modulation between active and sham tACS condition quantified via three modulation measures. Asterisks mark the uncorrected *p*-value of dependent samples t-tests. Only statistical tests for the first 1 sec time epoch after tACS offset remain significant after Bonferroni correction for comparisons in multiple time windows (α_bonf_ = 0.0167; 0.05/3). Error bars represent the standard error. * *p* < .05; ** *p* < .01.

### Spatial localization and dependency of tACS effects on individual physiology

The spatial specificity of tACS effects was analyzed in source space for the APL modulation parameter. Compared to D_KL_, APL is a more specific index assuming systematic amplitude modulations by tACS. Moreover, its computation is more stable compared to sine fit as no assumptions about parameter ranges are required. Cluster analysis in source space showed significantly increased APL values during active tACS relative to sham in a cluster over the parieto-occipital cortex (cluster test: *p* = .006) (Figure 4A). Significant differences were located primarily in the bilateral precuneus, superior parietal gyrus and midcingulate area as well as the cuneus and calcarine sulcus. These are part of flicker-entrained brain areas, located in tACS-targeted cortical regions and connected higher visual areas. Interestingly, tACS effects on SSR amplitude were not seen in the middle and superior occipital gyrus where neural phase alignment to the visual flicker peaked, but rather in neighboring occipital and parietal regions (insert in Figure 4B).

**Figure 4.**
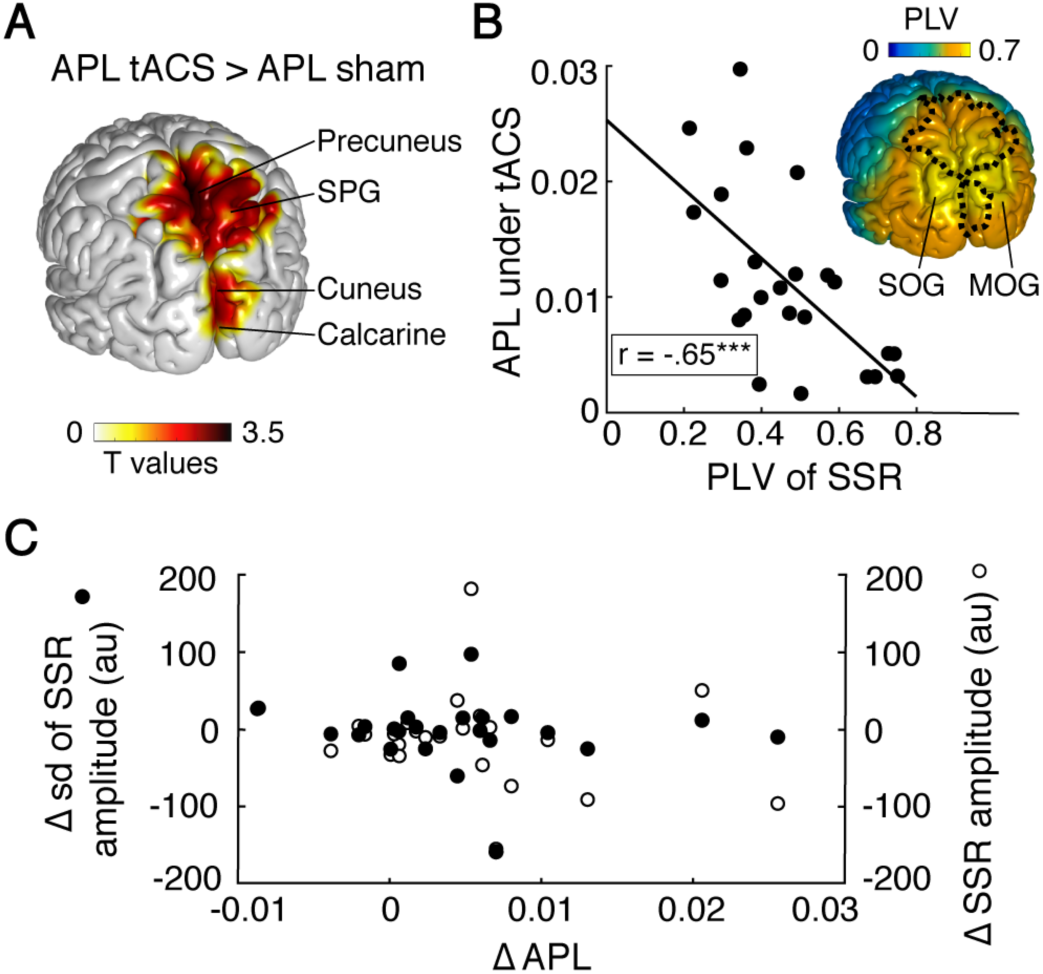
Analysis of tACS-induced amplitude phase locking (APL) of steady state responses (SSRs) during the first 1 sec epoch after tACS. (A) Significant differences in APL between active and sham tACS were localized in a cluster over the parieto-occipital cortex within flicker-entrained brain regions. (B) On sensor level, between-subject correlation shows that the tACS effect size quantified via APL is negatively related to EEG-flicker phase locking (PLV) during the tACS block (*r* = -.65, *p* < .001, n = 24). The insert illustrates that after source reconstruction, tACS effects (APL cluster boundary shown as dotted line) are mainly observed in regions where EEG-flicker PLVs did not peak during the tACS block. (C) The increase in APL values under tACS compared to sham is independent of overall changes in mean SSR amplitude or standard deviation (sd) of amplitude values over all trials per participant (sensor level data). MOG, middle occipital gyrus; SOG, superior occipital gyrus; SPG, superior parietal gyrus; *** *p* < .001.

**Figure 5.**
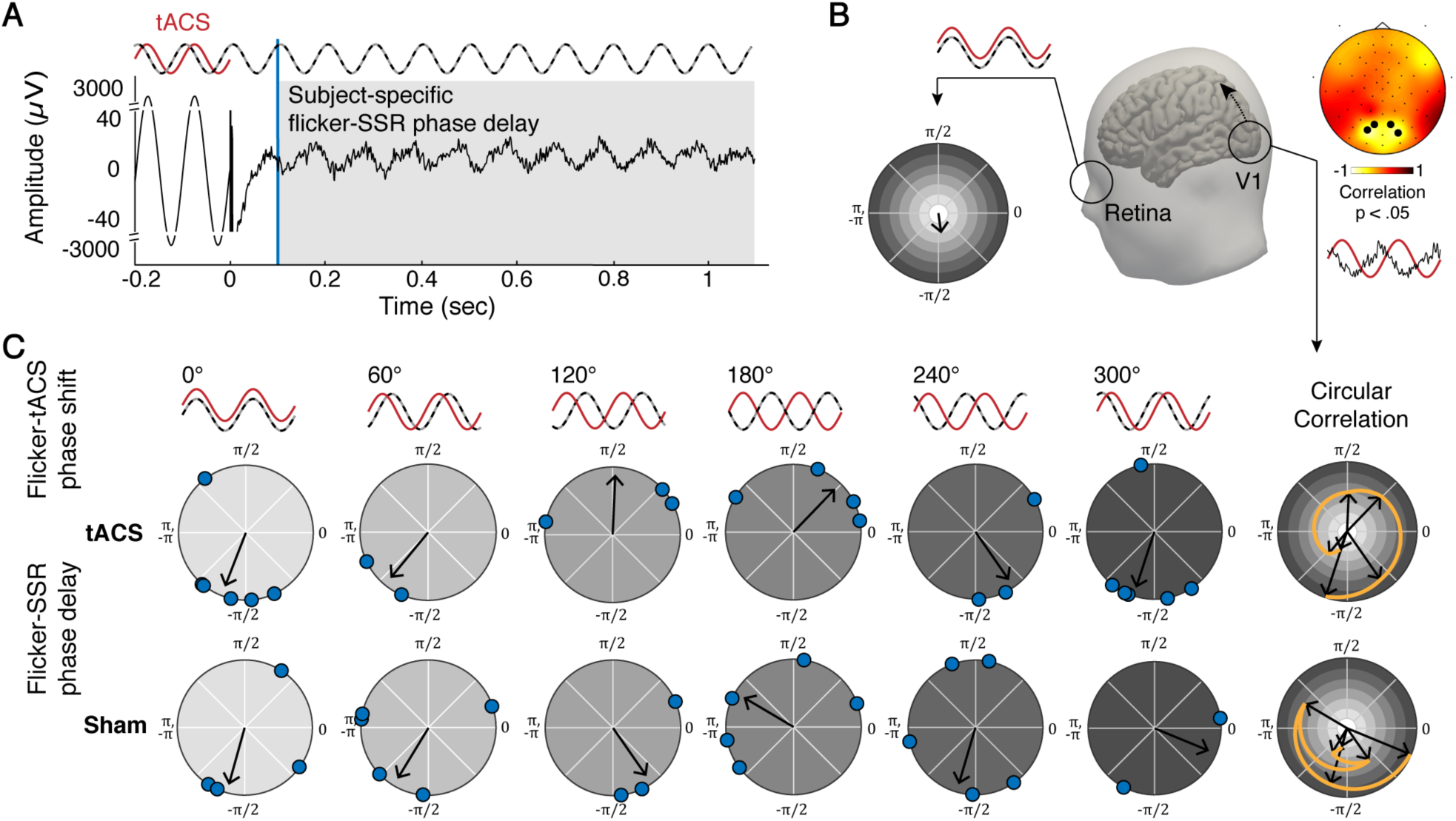
Analysis of the relation between individual cortical SSR onset delays and the optimal timing of tACS application associated with strongest SSRs. (A) Raw EEG data during the tACS block of one representative participant at electrode O1. Data during the last 0.2 sec of tACS stimulation show strong electrical artifacts. The gray-shaded area depicts the time window used for data analysis. (B) In case of an interference of flicker and electric field in the retina, visual response would be modulated before neural conduction delays along the visual pathway take effect. Thus, the optimal flicker-tACS phase shift should be independent of individual SSR onset delays and be comparable across participants as exemplified in the polar plot. In contrast, a significant cluster of occipital electrodes reveals a circular correlation between flicker-tACS phase shift and subject-specific SSR onset delays only during the tACS session. This suggests an interaction of tACS effects and SSR at early stages of visual cortical processing with concurrent spread of activity to higher visual areas. (C) Polar plots show flicker-SSR phase delays for all participants grouped by optimal flicker-tACS phase shift conditions exemplary for electrode O1. Blue circles show the mean flicker-SSR phase delay computed over all trials per participant and arrows depict the mean phase angle over all trials per optimal flicker-tACS phase shift condition. Circular plots on the right summarize the course of the mean flicker-SSR phase angles over the six flicker-tACS phase shift conditions (gray color code corresponds to flicker-tACS phase shifts of preceding polar plots). During tACS, data show a significant circular correlation between individual SSR phase delays and optimal flicker-tACS phase shifts across participants. There is no significant circular correlation during the sham session.

In accordance with this finding, we further observed a highly significant between-subject correlation of SSR amplitude modulation quantified by APL with the underlying strength of EEG-flicker PLV during the active tACS block on sensor level (*r* = -.65, *p* < .001). Participant with weaker PLV of SSR to flicker phase showed greater APL under tACS (Figure 4B). This relation was non-significant during the sham session (*r* = -.38, *p* = .067). As the difference in PLV between sham and tACS session showed no significant correlation with the difference in corresponding APL values (*r* = -.29, *p* = .166), data suggest that the strong negative relation between underlying PLV and SSR modulation is not generally valid but specific for tACS-induced neuromodulation. Our results thus emphasize the impact of the individual physiology, influenced by the neuronal response strength to visual flicker, for the detection of neural tACS effects.

Importantly, the increase in APL from sham to tACS could not be explained by a concurrent increase in mean SSR amplitude (*r* = -.22, *p* = .309) or standard deviation of SSR amplitude values (*r* = -.13, *p* = .556) over all trials of the tACS blocks per participant, thereby supporting the causal influence of the experimental manipulation of tACS-flicker phase shift (Figure 4C).

### Cortical level of interaction between SSRs and tACS effects

To examine the involvement of retinal co-stimulation on SSR amplitude modulation, we analyzed the dependency of optimal flicker-tACS phase shifts associated with strongest SSRs on individual flicker-SSR phase delays resulting from neural conduction times from the retina to visual cortex. Assuming an interaction between flicker and electric field at the retinal level before conduction delays of the visual system take effect, the SSR phase delay should have no predictive value for the optimal timing of tACS application. Notably, only during the tACS session, permutation tests revealed a significant cluster of occipital electrodes (O1, O2, O9, O10; cluster test: *p* = .009) showing a circular correlation between SSR phase delay and optimal flicker-tACS phase shift across participants. The mean correlation coefficient across cluster electrodes was *r* = -.54. Corresponding permutation tests during sham showed no significant electrode cluster.

## Discussion

Here, we used an innovative approach to study the phase-specificity of electrophysiological tACS effects in humans by systematically pairing rhythmic electrical with rhythmic visual stimulation. Specifically, we evaluated the amplitude of flicker-evoked SSRs in the interval immediately following tACS delivered at six different phase shifts relative to flicker onset. We demonstrate that SSR amplitudes are modulated in a tACS phase-dependent manner and that tACS effect size is inversely related to the strength of the underlying flicker response. Furthermore, we found that the individual phase delay between flicker onset and cortical SSR is predictive of the optimal timing of tACS application.

Previous investigations into the tACS mechanism of action typically aimed to modulate ongoing or task-related brain oscillations which show high variability in frequency and phase consistency. Here, we used SSRs that allow the precise setting of the oscillatory phase in the visual cortex. While the neural mechanism underlying SSR generation is still debated, increasing evidence supports the involvement of entrained endogenous oscillations in the processing of rhythmic visual input [41–43]. Yet, irrespective of their exact physiological origin, SSRs can be treated as stimulus-driven brain rhythms with predictable time course, which enables a highly specific targeting by tACS. Notably, compared to ongoing activity, the phase of the SSR is not expected to be shifted by tACS as EEG recordings will always show dominant phase locking to the driving flicker. Rather, if voltage modulations by electrical and sensory stimulation interact in a phase-specific manner, SSRs should reflect this interaction in net amplitude modulations. Our data confirmed that SSR amplitudes measured immediately after tACS offset are dependent on the relative phase shift between flicker and tACS which provides first conclusive evidence for the phase-specificity of tACS effects in human EEG recordings. Importantly, as data show no overall amplitude increase of SSRs by tACS (Figure 3D), results imply a phase-dependent enhancement and suppression of flicker responses due to the sinusoidal nature of the tACS signal. This is in accordance with recent invasive recordings in animals, showing that tACS is capable of modulating the timing of neural spike discharges [7,8].

The phase-dependent modulatory effect on SSRs was robust under three modulation measures but already decayed after the first 1 sec time epoch. While lasting power enhancement after prolonged alpha tACS has been repeatedly shown and ascribed to synaptic plasticity mechanisms [30,31,44], evidence for outlasting effects on phase synchronization is still sparse. Yet, neural oscillatory activity entrained during rhythmic electric, magnetic or visual stimulation were shown to remain stable for a few cycles after stimulation offset [45–47]. These short-lived reverberations of oscillatory activity, phase-locked to the external force, have been termed entrainment echoes [36,46]. Notably, Vossen et al. [30] could not find evidence for entrainment echoes after short tACS epochs of 3 or 8 sec length. However, while tACS phase effects might not be detectable via PLV in ongoing brain activity, flicker-induced brain rhythms may increase the signal-to-noise ratio and thereby enhance the detectability of short-lived tACS aftereffects. These results are compatible with the assumption of online tACS-induced entrainment of neural activity, which yet cannot be tested directly due to artifact-contaminated EEG recordings. We speculate that the observed effects generalize to tACS and SSRs in other frequency bands of the human EEG, which should be systematically tested in future studies.

Interestingly, the efficacy of tACS was inversely related to the strength of the underlying flicker response. More precisely, tACS-induced SSR amplitude modulation was stronger in participants showing weaker phase locking between flicker and evoked oscillatory activity. Furthermore, tACS effects were especially prominent in brain areas that were not maximally driven by the driving flicker. While maximal phase locking to the visual stimulus occurred in the middle and superior occipital gyrus, maximal SSR amplitude effects of tACS were seen in early visual cortex and superior parietal cortex. As tACS effects were also observed in downstream visual cortical areas, these results suggest that stimulation effects might not only act locally in regions targeted by the electric field but also affect connected areas via polysynaptic interactions [48–50]. Moreover, these findings emphasize a dependency of tACS effects on the individual physiology. It has repeatedly been demonstrated that tACS effect sizes are related to the baseline level of oscillatory activity, with effects only being measurable when power at the stimulated frequency was low [51–53]. Similarly, our data indicate that in participants and anatomical regions exhibiting strong EEG-flicker phase locking, the dominance of synchronous neural activity may have led to ceiling effects in alpha amplitude with no room for further enhancement. Also, the ratio between the strength of the external tACS stimulation relative to the strength of the neural oscillation was suggested as a critical factor for successful entrainment [53], that may explain differential tACS effect sizes for varying flicker-evoked oscillation strengths.

Important to consider in the interpretation of tACS effects is the possible involvement of sensory co-stimulation effects, that may indirectly lead to neural entrainment. First, electric stimulation of peripheral nerves in the skin can induce rhythmic activation of the somatosensory cortex that was shown to at least partly explain tACS effects on motor cortex [27]. Yet, recent studies demonstrated that tACS affects neural activity even if somatosensory input is controlled for [7,54]. In our study, the dependency of tACS effects on the SSR phase delay over the primary visual cortex, i.e., the target region of the electrical stimulation, point to transcranial stimulation effects on SSR at early stages of visual cortical processing with concurrent spread of activity to higher visual areas. However, to rule out entirely that SSRs were influenced by somatosensory co-stimulation, a control electrode montage would be necessary that elicits equivalent tactile sensation without stimulating the cortex. This is challenging as cutaneous stimulation effects depend not only on current strength but also on the distribution and sensitivity of skin receptors in the stimulated area [55]. Therefore, the development of valid control montages is one of the key goals in future tACS research. Second, it has been noted that tACS can cause activation of the retinae due to shunting of scalp-applied currents across the skin. Although no participant reported phosphene sensation with the used focal ring montage, sub-threshold activation of the retinae could in principle have an impact on visual processing, which was previously dealt with by using control electrode montages [55]. Here, our experimental approach allowed to directly disentangle cortical from retinal stimulation effects. It is well known that flicker stimulation will inevitably lead to variable delays in the recordable SSR that likely result from interindividual differences of the conduction times along the visual pathway. Accordingly, a cortical interaction between tACS effects and SSRs should be associated with varying optimal phase shifts between tACS and flicker across participants. This was exemplified in a study by Ruhnau et al. [56], showing that simultaneously started tACS and flicker did not lead to tACS-induced power increases of the fundamental SSR component in the overall sample. In our study, we could show that the optimal flicker-tACS phase shift associated with strongest SSRs was correlated with the individual flicker-SSR phase delay over the primary visual cortex. This finding suggests that it is possible to predict the optimal timing of tACS application based on subject-specific conduction delays of the visual system. Importantly, this dependency cannot be explained by an interference between flicker and electric field at the retinal level, which should be independent of postretinal neural conduction times and, therefore, result in comparable optimal flicker-tACS phase shifts across participants. Thus, our finding strongly suggests that the interaction between tACS effect and SSR was cortical rather than retinal in nature.

In conclusion, we provide the first conclusive electrophysiological evidence for the efficacy of tACS to modulate cortical oscillatory activity in a phase-dependent manner in humans. Importantly, we found that the optimal phase shift between electrical and visual stimulation varied between participants and depended on baseline conduction delays of the visual system. This result cannot be explained by peripheral electric co-stimulation of the retina and thus strongly supports a cortical origin of tACS effects. Furthermore, our data corroborate the notion that tACS effect size is limited by the strength of the targeted oscillatory activity level. Taken together, our findings show that individual functional properties and the current oscillatory brain state need to be considered to achieve specific, custom-fit neuromodulation. In the context of the ongoing debate on the overall capability of tACS to entrain oscillatory activity in humans, our results make a strong case for its periodic impact on neuronal excitability that would allow for frequency- and phase-specific modulations of brain signals.

## Acknowledgements

This work was supported by the Deutsche Forschungsgemeinschaft (SFB 936/A3 awarded to A.K.E. and T.R.S.; SPP 1665/EN 533/13-1 and SFB TRR 169/B1 awarded to A.K.E.; SPP 1665/SCHN 1511/1-2 awarded to T.R.S.). We thank Jonathan Daume, Alexander Maye, Jan-Ole Radecke and Darius Zokai for helpful discussions on the data, Malte Sengelmann and Guido Nolte for methodological support, and Karin Deazle and Darius Zokai for assistance in data recording.

